# At the Intersection Between SARS-CoV-2, Macrophages and the Adaptive Immune Response: A Key Role for Antibody-Dependent Pathogenesis But Not Enhancement of Infection in COVID-19

**DOI:** 10.1101/2021.02.22.432407

**Authors:** Jennifer K. DeMarco, Wiliam E. Severson, Daniel R. DeMarco, Gregory Pogue, Jon Gabbard, Kenneth E. Palmer

## Abstract

Since entering the world stage in December of 2019, SARS-CoV-2 has impacted every corner of the globe with over 1.48 million deaths and caused untold economic damage. Infections in humans range from asymptomatic to severe disease associated with dysregulation of the immune system leading to the development of acute respiratory distress syndrome (ARDs).

The distinct shift in peripheral monocyte activation and infiltration of these cells into the respiratory tract in ARDs patients suggests severe COVID-19 may largely result from damage to the respiratory epithelia by improperly activated macrophages. Here, we present evidence that dysregulation of the immune response in COVID-19 begins with activation of macrophages by non-neutralizing antibodies and induction of ACE2 expression, rendering these cells susceptible to killing by SARS-CoV-2. Death of macrophages occurs independently of viral replication and leads to the release of inflammatory mediators and modulation of the susceptibility of downstream epithelial cells to SARS-CoV-2.

## Introduction

Following the appearance of COVID-19 in December 2019 in Wuhan, China, SARS-CoV-2 has swept across the globe at a rate not seen since the Spanish influenza of 1918 (1-3). A high rate of transmission combined with a large percentage of asymptomatic/mild carriers has led to more than 62 million cases and 1.48 million deaths, as well as a devastating economic toll worldwide (1-3). Following its emergence, SARS-CoV-2 was rapidly identified by Zhu et al (1) as a betacoronavirus closely related to SARS-CoV and MERS-CoV. Like other members of this family, SARS-CoV-2 has a large genome of approximately 30Kbp surrounded by an envelope comprised of S (spike), N (nucleocapsid), E (envelope), and M (membrane) proteins (3). Of these, the spike protein, a type I trimeric glycoprotein, has received the most attention as a target for both potential vaccine and therapeutic development due to its involvement in host receptor binding and viral uptake (3). The spike protein harbors two primary regions: an S1 domain containing a receptor binding region (RBD) able to bind to the human angiotensin II converting enzyme receptor (ACE2) and an S2 domain containing a furin cleavage site that facilitates fusion of the virus and host cell membrane following receptor binding (3). Studies with SARS-CoV and MERS-CoV have suggested that neutralization of the receptor binding domain (RBD) may provide protection from infection, making this region a key target for vaccine and therapeutic development (3).

While the majority of patients with SARS-CoV-2 present with mild to moderate respiratory symptoms, approximately 15% develop severe disease with a high rate of mortality due to ARDs (4). Although risk factors influencing expression of ACE2 (diabetes, high blood pressure, kidney and cardiovascular disease) are now well documented, the factors that determine the outcome of COVID-19 cases remain elusive (4-5). Furthermore, it is now known that a subset of patients with mild to moderate cases will also go on to develop long-term symptoms of COVID-19, though whether this is the result of direct tissue damage by the virus or ongoing inflammatory responses is not yet clear (4-6).

Recent clinical studies have characterized two distinct phases of COVID-19 (4-6): (1) an early phase with relatively mild symptoms during the first five days of infection with high levels of viral occurring in susceptible tissues and (2) a late phase in which severe symptoms including ARDs, hypercoagulability and multi-organ failure emerge as a result of immune dysregulation and associated tissue damage (4-6). The delay in onset of disease severity and correlation to the emergence of the first antibodies, suggests that induction of the adaptive immune response and the antibody-antigen presenting cell (APC) interface may be in part responsible for the majority of morbidity and mortality associated with COVID-19 (4-7). Comparison of hematological profiles between patients with relatively mild disease and those with ARDs supports this hypothesis, showing marked elevation in CD14+ IL-1b producing monocytes in the peripheral circulation (7), infiltration of monocytes and neutrophils into the respiratory tract (4)as well as IL-6 and IL-1b in the serum of ARDs patients (4,7). Taken together with the finding that multiple serological markers in severe COVID-19 patients that mirror those observed in macrophage activation syndrome (MAS) such as hyperferritinaemia, altered liver function and coagulopathy, suggests a critical role for macrophages in determining the outcome of SARS-CoV-2 infection (4).

To date, there has been considerable controversy surrounding the role of APCs in COVID-19 and whether they are susceptible (8-9) or recalcitrant (10-12) to SARS-CoV-2 infection. Expression of ACE2 only occurs on a subset of CD14+ monocytes (11-12) and given the relationship between inflammation and ACE2 expression in epithelial cells (13), caution is in order when interpreting these results as many of the techniques used to culture macrophages may unintentionally polarize these cells towards an activated phenotype. Monocytes and macrophages are first on the scene in response to pathogens and responsible for many components of the immune response to novel pathogens including recruitment of T cells and neutrophils to the site of infection, presentation of bound antigen-antibody complexes thru Fc receptors, secretion of inflammatory mediators to support lymphocyte development and initiation of wound healing responses post-infection (8-12). Given the correlation in the timing of the emergence of antibodies with severe COVID-19 and the similarity of late stage COVID-19 to MAS (4, 8-12), we examined the role of antibodies in modulating the inflammatory response in murine macrophages and show that non-neutralizing antibodies to SARS-CoV-2 induce ACE2 expression in murine macrophages, rendering them susceptible to replication-independent killing by SARS-CoV-2. Supernatants from these cells exhibit a cytokine profile similar to those observed in the serum of ARDs patients and addition of these supernatants to Vero E6 cultures significantly enhances susceptibility to SARS-CoV-2. These results suggest a model for antibody-dependent induction of immune dysregulation in COVID-19.

## Results

Given the expression of ACE2 in only a small subset of peripherally derived CD14+ cells (11-12), we asked whether activation of ACE2-negative macrophages could induce ACE2 expression and render macrophages susceptible to SARS-CoV-2. No reduction in viability was observed between naive macrophages and those incubated with SARS-CoV-2 in the absence of antibody. Addition of non-neutralizing antibodies to either the intracellular nucleocapsid antigen or surface-expressed spike protein reduced cell viability to 35.98% (p<0.0001) and 53.67% (p<0.0001), respectively (Figure 1). By stark contrast, sensitization to viral killing by SARS-CoV-2 did not occur in the presence of neutralizing antibodies to the receptor-binding domain (RBD), suggesting that antibody binding to macrophages may be able to induce ACE2 expression and that availability of the RBD domain is required for virus-induced cytotoxicity. Subsequent analysis by qPCR revealed no increase in viral load, indicating that death is not due to lytic replication (Figure 1).

**Figure 1:**
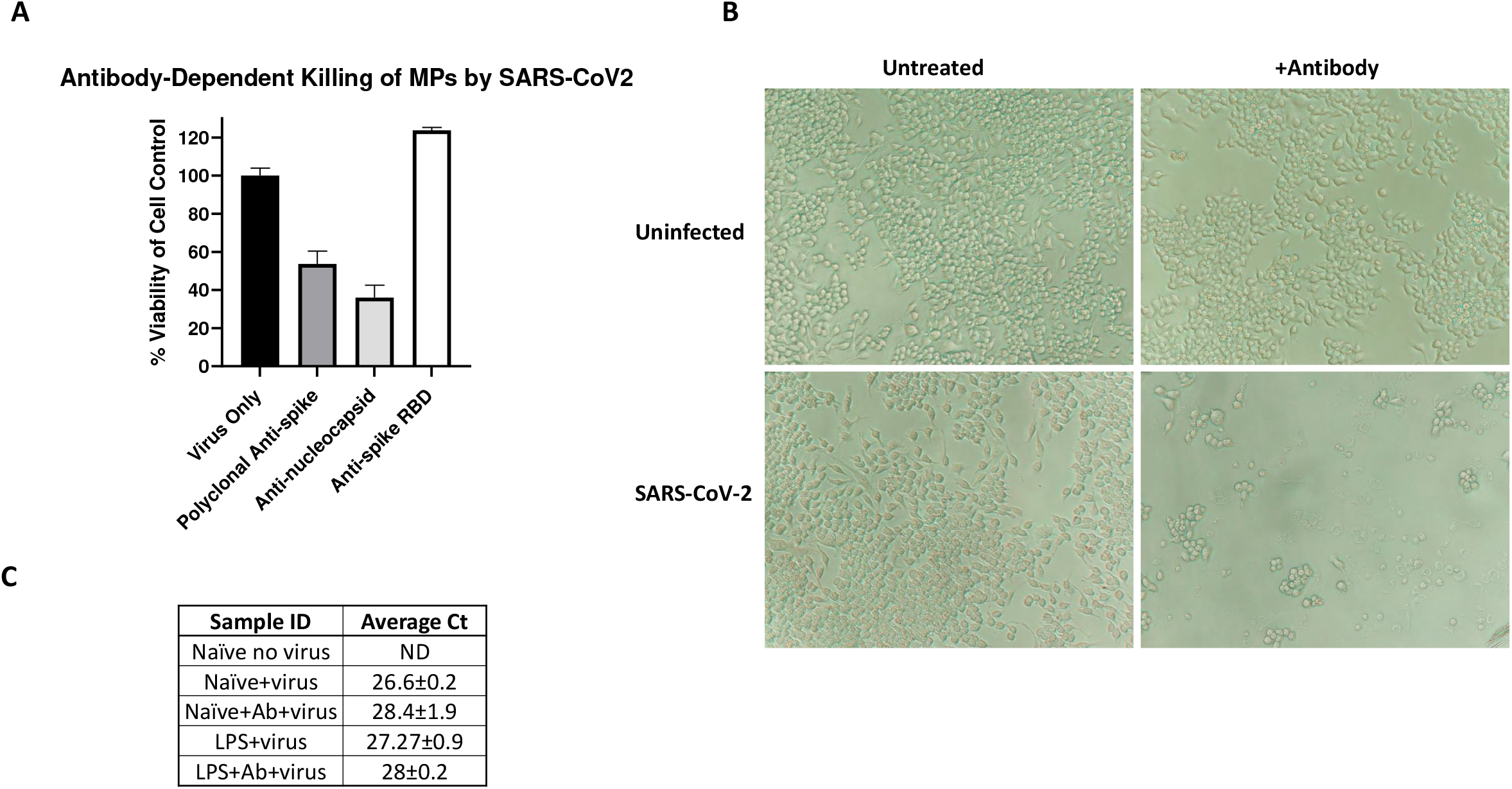
Antibody-dependent killing of macrophages by SARS-CoV-2. A-B) In the absence of antibody, Raw 264.7 cells are resistant to killing by SARS-CoV-2. Addition of non-neutralizing antibodies against either the nucleocapsid or spike protein reduced survival at 48 hours post-infection to 35.98% and 53.67% of the cell control (p<0.0001). Neutralization of the RBD prevents viral killing, suggesting that binding of ACE2 may be required for antibody-dependent infection of monocytes and macrophages. C) No increase in virus load was observed between naïve and activated macrophages incubated with SARS-CoV-2, suggesting that death is not due to lytic replication.

To confirm that antibody-induced susceptibility to SARS-CoV-2 resulted from induction of ACE2 expression following Fc ligation, we evaluated expression of ACE2 in Raw264.7 cells incubated with virus in the presence or absence of antibodies or LPS by fluorescent microscopy. In the absence of non-neutralizing antibodies, macrophages were found to express only minimal levels of ACE2 (Figure 2). Following activation either by antibody-dependent Fc ligation or Toll-like receptor 4 (TLR4) induction after exposure to LPS expression of ACE2 increased by 49.5 and 48.9-fold respectively (p<0.0001, Figure 2).

**Figure 2:**
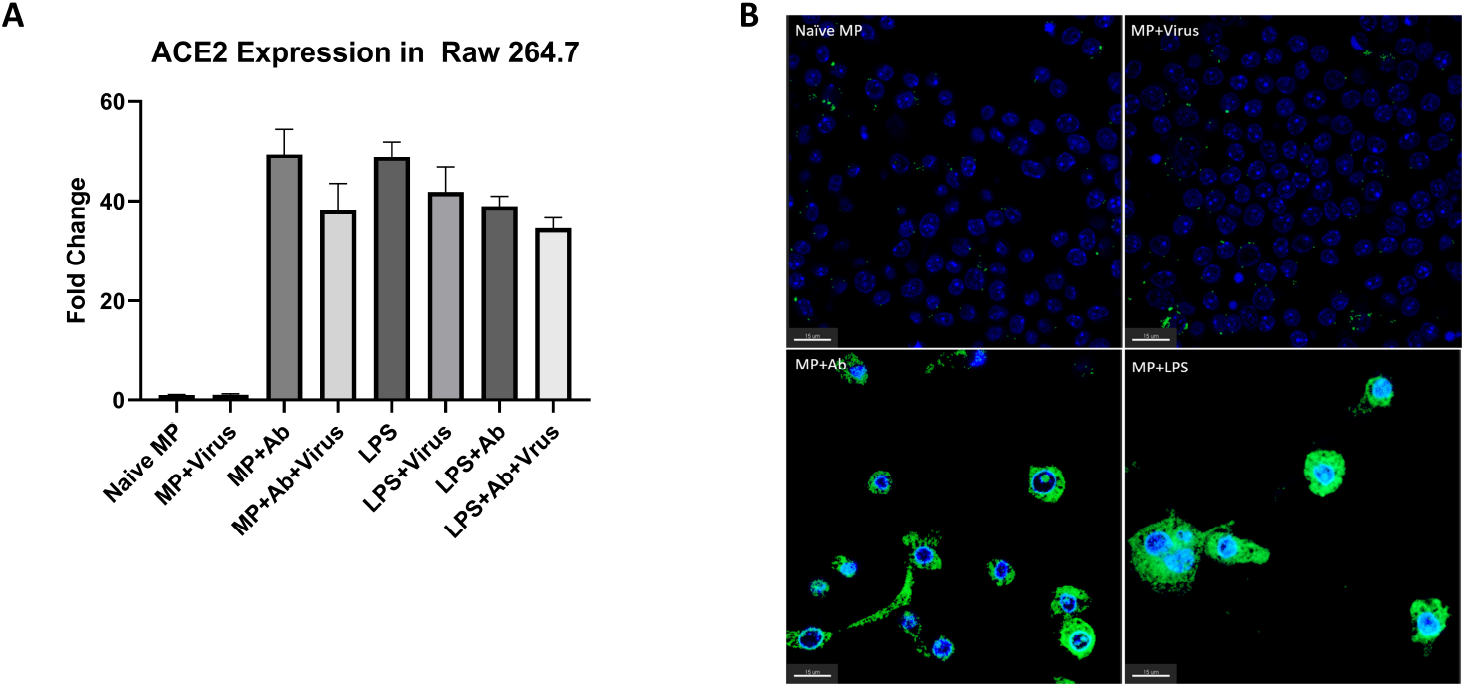
Activation of Raw264.7 cells induces ACE2 expression and susceptibility to SARS-CoV-2. Following incubation with polyclonal anti-spike or LPS and/or SARS-CoV-2 as described above, Raw264.7 cell were fixed and evaluated for ACE2 expression via confocal microscopy. A.) Fold changes were calculated as a ratio of the experimental group: naïve macrophages. Activation of Raw264.7 cells by Fc ligation or through LPS exposure induced expression of ACE2 nearly 48-fold (p<0.0001) and B.) resulted in morphological changes consistent with macrophage activation. Blue, DAPI; green, ACE2.

Given the finding that treatment with LPS alone induced ACE2 expression, we examined the susceptibility of LPS-activated macrophages to SARS-CoV-2. Exposure to LPS prior to virus challenge reduced macrophage survival following virus challenge to 77.35% in the absence of antibody (p<0.0149, Figure 3). Addition of non-neutralizing antibody further reduced survival to 11.13%, significantly less than that observed in the presence of antibody alone (44.94%, p<0.0001; Figure 3). These data suggest that induction of ACE2 expression is only one component required for sensitization of macrophages to SARS-CoV-2 and that Fc ligation by non-neutralized antibody-virus complexes are a key component of this process.

**Figure 3:**
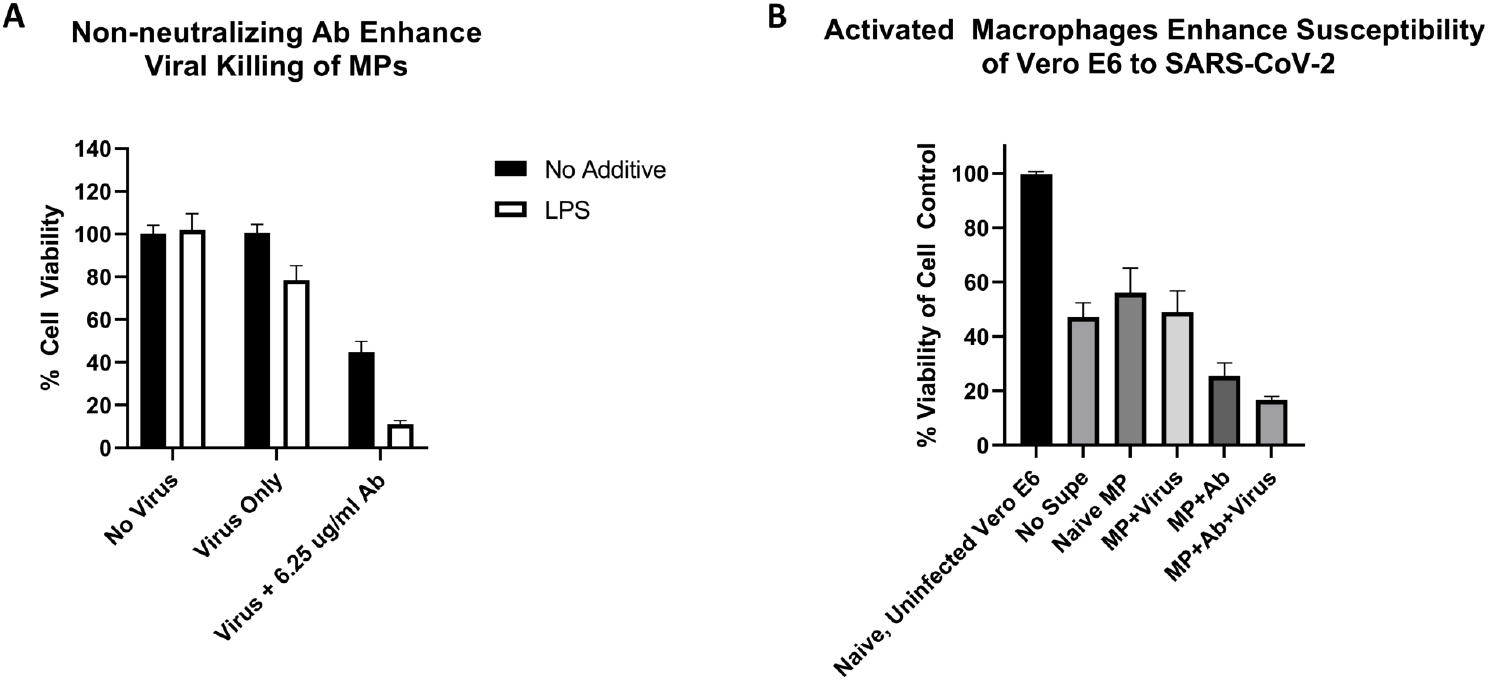
Activation of Raw264.7 cells thru TLR4 enhances susceptibility to SARS-CoV-2. A) Activation of RAW264.7 cells with LPS enhanced the susceptibility of macrophages in the absence of antibody, reducing survival to 77.35% after 48 hours of incubation with SARS-CoV-2 (p=0.0149). Addition of non-neutralizing antibodies further reduced survival to 11.13%, significantly less than that observed in the presence of antibody alone (44.94%, p<0.0001). **B)** The addition of supernatants from SARS-CoV-2 infected macrophages enhances viral killing of Vero E6 cells by nearly 2.8-fold (p=0.011).

To further characterize the potential impact of antibody-dependent susceptibility of macrophages to SARS-CoV-2 on downstream cell populations, we next examined the impact of macrophage supernatants on susceptibility of Vero E6 cells to SARS-CoV-2. Supernatants from macrophages exposed to SARS-CoV-2 had no significant impact on susceptibility of Vero E6 to SARS-CoV-2 (Figure 3). However, those from macrophages activated with antibodies in the presence of SARS-CoV-2 enhanced viral killing of Vero E6 cells by 2.8-fold (p=0.011). This suggests that macrophages infected with SARS-CoV-2 by antibody-based induction of susceptibility to SARS-CoV-2 may be directly responsible for enhancing viral damage to respiratory epithelial cells in severe COVID-19.

From a comparison of recent clinical studies, elevation of sixteen cytokines (TNF-a, IL-6, IL-10, RANTES, IL-1b, IL-2, GM-CSF, IL-18, IP-10, IL-4, IFN-y, IL-9, G-CSF, MCP-1, IL-17a and MIP-1a) have emerged as hallmarks of severe COVID-19 (14). Of these, we observed that nine (TNF-a, RANTES, IL-6, IL-1b, GM-CSF, IL-18, IFN-y, G-CSF and MIP-1a) were found to be reduced in the supernatants of macrophages activated through Fc ligation and markedly increased after these cells were infected with SARS-CoV-2 (Figure 4 and Table 1). Five additional cytokines (IL-2, IP-10, IL-4, IL-9 and IL-17a) were induced only with SARS-CoV-2 infection (See Figure 5), suggesting that induction of pro-inflammatory mediators in APCs likely occurs through at least two pathways: one that is Fc-dependent and one that results directly as a result of SARS-CoV-2 infection. Of note, IL-10 was only elevated in the supernatants of LPS-activated suggesting that either macrophages are not directly responsible for the elevated levels of IL-10 observed in COVID-19 patients or induction of IL-10 occurs in the final stages of the cytokine storm through activation of additional inflammatory signaling pathways.

**Figure 4 :**
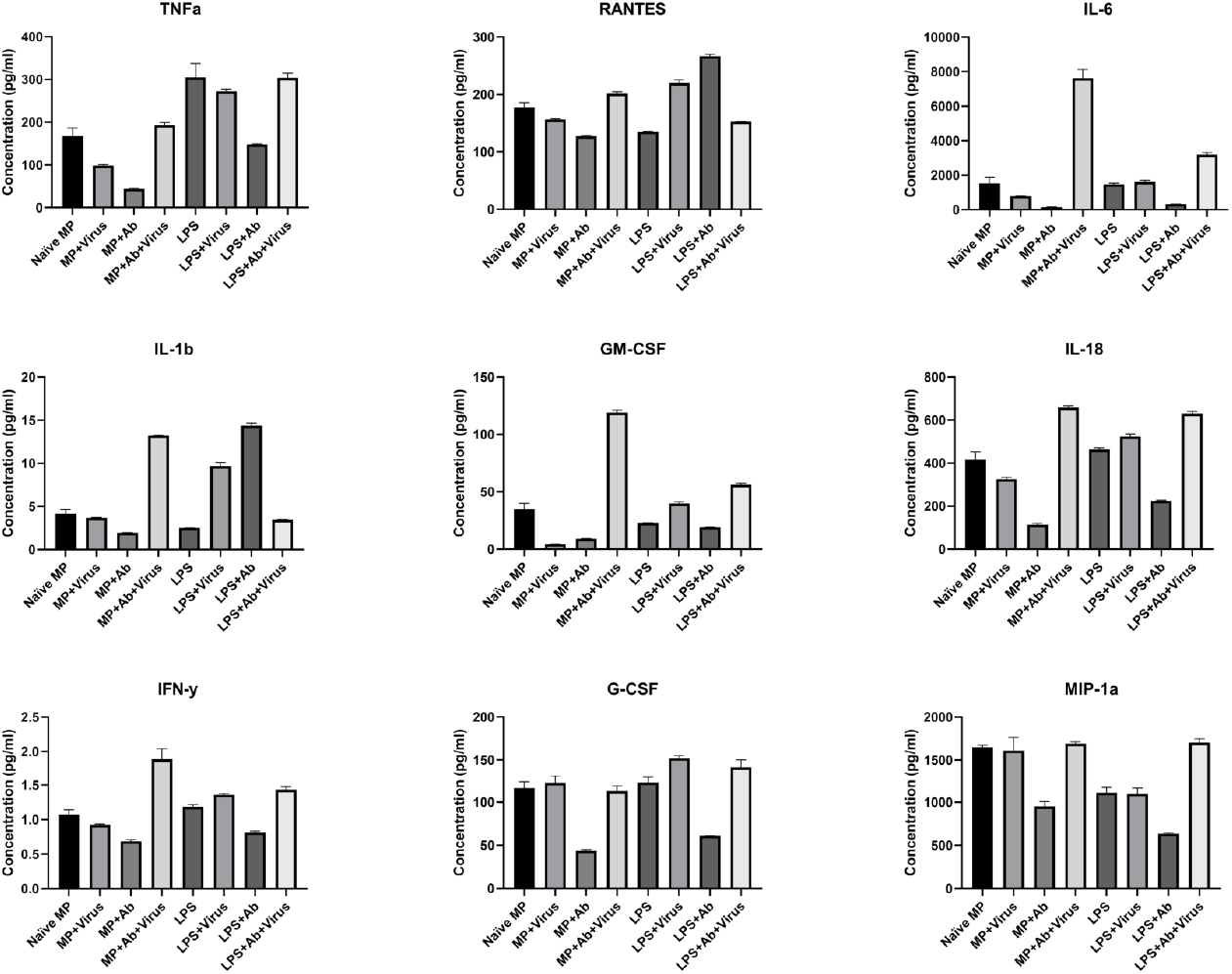
Modulation of inflammatory mediators released from SARS-CoV-2 occurs thru an antibody-dependent process. The release of nine cytokines (highlighted) identified as hallmarks of severe COVID-19 infection by Wang et al^12^ from macrophages were modulated by Fc ligation alone (MP±Ab). Subsequent infection with SARS-CoV-2 increased production of all nine, as well as IL-2, IP-10, IL-4, IL-9 and MIP-1a suggesting that at least two distinct pathways are activated in response to infection with SARS-CoV-2 in macrophages (MP+Ab±Virus). Significant induction of IL-10 was only observed in macrophages activated via TLR4 induction (MP+Ab+Virus±LPS).

**Table 1 :** Modulation of inflammatory mediators released from SARS-CoV-2 occurs thru an antibody-dependent process. The release of nine cytokines (highlighted) identified as hallmarks of severe COVID-19 infection by Wang et al^12^ from macrophages were modulated by Fc ligation alone (MP±Ab). Subsequent infection with SARS-CoV-2 increased production of all nine, as well as IL-2, IP-10, IL-4, IL-9 and MIP-1a suggesting that at least two distinct pathways are activated in response to infection with SARS-CoV-2 in macrophages (MP+Ab±Virus). Significant induction of IL-10 was only observed in macrophages activated via TLR4 induction (MP+Ab+Virus±LPS).

|  | MP±Ab | MP+Ab±Virus | MP+Ab+Virus±LPS |
| --- | --- | --- | --- |
| **TNF-a** | 0.0003 | <0.0001 | 0.0012 |
| **RANTES** | <0.0001 | <0.0001 | NS |
| **IL-6** | 0.0089 | <0.0001 | <0.0001 |
| **IL-10** | NS | NS | <0.0001 |
| **IL-1b** | 0.0002 | <0.0001 | <0.0001 |
| **IL-2** | NS | 0.0006 | <0.0001 |
| **GM-CSF** | <0.0001 | <0.0001 | <0.0001 |
| **IL-18** | <0.0001 | <0.0001 | NS |
| **IP-10** | NS | <0.0001 | <0.0001 |
| **IL-4** | NS | <0.0001 | 0.0013 |
| **IFN-γ** | 0.0141 | <0.0001 | 0.0031 |
| **IL-9** | NS | 0.0343 | NS |
| **G-CSF** | <0.0001 | <0.0001 | NS |
| **MCP-1/CCL-2** | NS | NS | NS |
| **IL-17a** | NS | 0.0034 | 0.001 |
| **MIP-1a/CCL-3** | <0.0001 | <0.0001 | NS |

**Figure 5:**
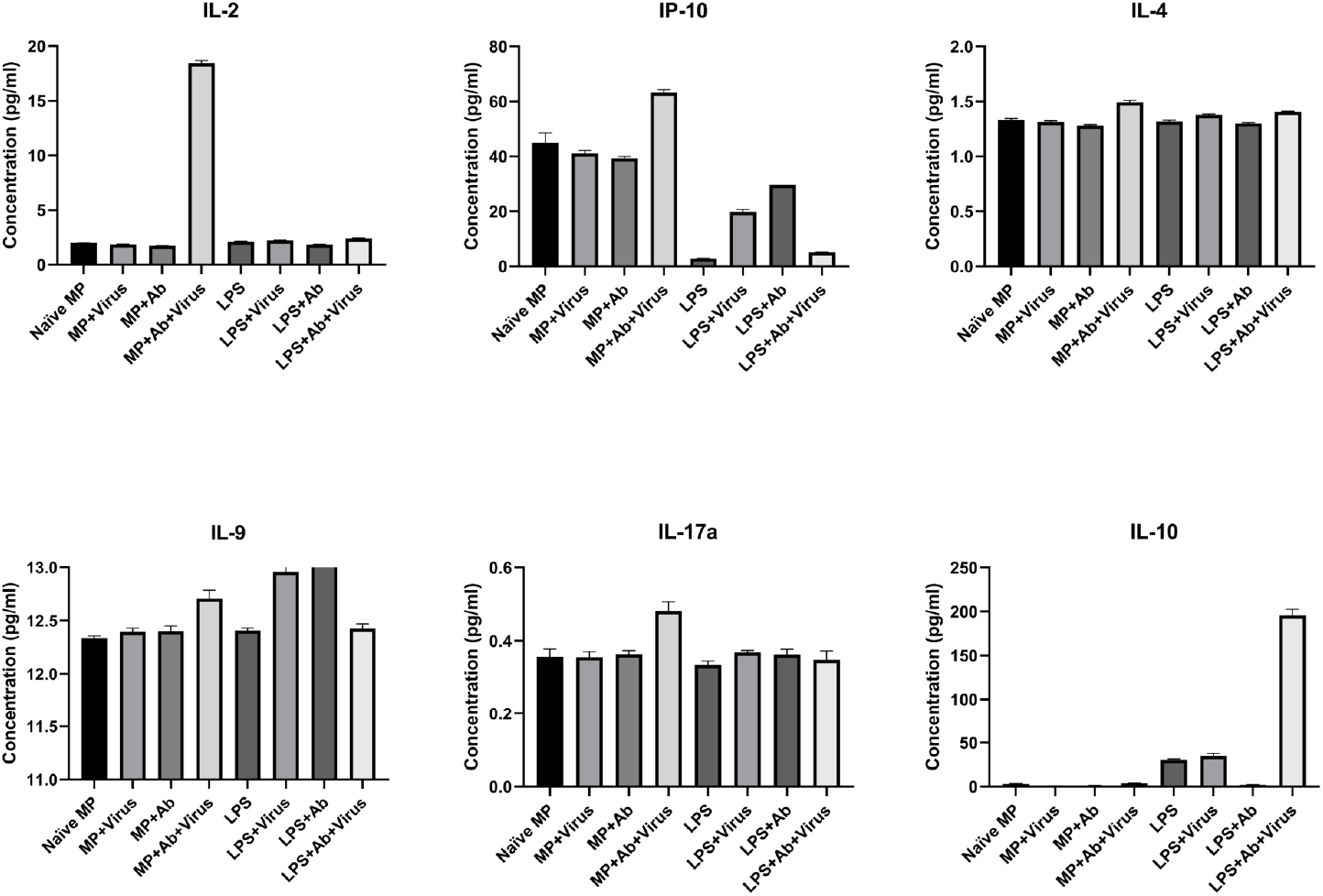
Evidence for multiple pathways for macrophage activation in SARS-CoV-2 infection. Five additional markers, IL-2, IP-10, IL-4, IL-9 and IL-17a were observed to be modulated only after infection of macrophages with SARS-CoV-2, suggesting that there are at least two separate pathways (Fc-dependent and Fc-independent) through which SARS-CoV-2 initiates production of cytokines in macrophages. Induction of IL-10 was only observed in LPS-activated macrophages suggesting that either macrophages are not directly responsible for the increase in IL-10 observed in ARDs patients or that secondary activation of macrophages through Toll-like receptors may play a role during the later stages of the disease.

## Discussion

The high degree of correlation observed between migration of mononuclear cells into infected tissue and a pattern of MAS-like inflammatory markers that occur in patients with severe disease has led to the morbidity and mortality in COVID-19 being attributed to dysregulation of macrophages in the later stages of the infection (15-16). The onset of ARDs coincides with the emergence of antibodies at the transition between the innate and adaptive immune response, suggesting that immune modulation by newly circulating antibodies may be important to initiating a hyperinflammatory state in susceptible individuals. The timing of onset of severe symptoms also argues that regardless of the initiating factor(s), the pathway leading to immune dysregulation in COVID-19 is not an innate state but must be induced during development of the adaptive immune response. This is notable, particularly in the context of APCs as it suggests that the populations key to understanding the pathogenesis of COVID-19 are those that are initially refractory to infection with SARS-CoV-2 and that contribution of the subset of CD14+ cells that innately express ACE2 are likely minor in comparison to those that come into play as antibodies emerge. We observed that naïve macrophages express little ACE2 on the cell surface and are resistant to killing by SARS-CoV-2. Non-neutralizing antibodies render these cells susceptible via induction of ACE2 expression, suggesting that the cascade leading to ARDs in COVID-19 may begin with the ligation of Fc receptors on APCs at the onset of the adaptive immune response.

The ability of TLR induction to induce ACE2 expression and render macrophages susceptible to SARS-CoV-2 is notable as a number of viruses promote type I interferon signaling through induction of the TLR4/MyD88 axis (17). A recent study by Duan et al (16) characterizing the role of macrophage polarization demonstrated that although M1 and M2 macrophages show similar competency to eliminate SARS-CoV-2, M1 polarized macrophages contributed to respiratory damage whereas M2 polarized macrophages cleared the virus without causing inflammatory-medicated injury. Given that M1 and M2 polarization are typically exclusive states, the high level of ACE2 receptor induction in LPS-activated (M1) polarized macrophages may be something that does not occur in M2 polarized cells. This warrants further investigation, and if confirmed, may suggest that inappropriate polarization of macrophages to an M1 phenotype, whether pre-existing or induced by SARS-CoV-2 infection, sets the stage for ARDs.

It is important to note that induction of ACE2 receptor expression only constitutes part of the story as addition of non-neutralizing antibodies enhanced viral killing of LPS-activated macrophages by nearly 2-fold that observed with LPS alone. This suggests that not only is induction of ACE2 receptor expression required to render macrophages susceptible to SARS-CoV-2, but that binding of virus by antibodies may also enhance uptake of receptor-bound virus.

Antibody-dependent enhancement has been a point of concern in the rapid development of vaccines for COVID-19 as ADE has been documented with other betacoronaviruses in the experimental setting and has thwarted development vaccines for feline coronaviruses, as well as a number of others for decades (18). Dengue fever is perhaps the best-known model of ADE. Infection of naive hosts with dengue virus results in a classical presentation of DF, which is typically self-limited in immunocompetent hosts (18-19). Subsequent infection with a strain different carries a significant risk of developing dengue hemorrhagic fever (DHF) as a direct result of harboring pre-existing antibodies that not only fail to neutralize the second virus, but also enhance viral uptake and replication (18-19). Compared to DF, DHF carries a significantly higher morbidity and mortality rate (18-19).

Models of ADE developed from the classical presentation of enhanced disease following re-infection in the Flaviridae (18-19), form the basis of the current ADE paradigm, wherein non-neutralizing antibodies, either from vaccination or previous infection enhance two key components of viral pathogenesis: 1) viral uptake into normally non-permissive cell populations and 2) subsequent enhancement of viral replication. COVID-19 is difficult to reconcile with the established paradigm for ADE as SARS-CoV-2 is able to infect and kill, but not replicate in several key leukocyte populations (8-12). However, as a direct result of abortive replication, SARS-CoV-2 induces a pyroptosis-like cell death that results the release of inflammatory mediators into the extracellular space (6, 9, 20). Our findings that supernatants from SARS-CoV-2 infected macrophages enhance susceptibility in Vero E6 cells and correlate with published profiles of cytokine and chemokine expression in COVID-19 patients (4, 13, 21), suggests the need for an updated model of antibody-dependent pathogenesis in COVID-19 wherein non-neutralizing antibodies activate and render APCs susceptible to SARS-CoV-2. Death of infected APCs releases cytokines and chemokines which are able to directly alter the outcome of infection in epithelial cells and initiate a cascade of proinflammatory stimuli leading to a multi-component cytokine storm. This model would account for the limited efficacy observed in clinical trials of therapeutic agents targeting individual cytokines, such as tociluzimab (anti-IL-6, 22-23) and TNF-α^6^ compared to broad-spectrum inhibition of the inflammatory response with dexamethasone (24).

The importance of the macrophage and non-neutralizing antibodies at the intersection of the innate and adaptive immune response in COVID-19 has significant ramifications for the use of convalescent plasma and the development of effective vaccines. The antibodies that comprise convalescent plasma vary greatly from donor to donor, in both the ratio and affinity of neutralizing versus non-neutralizing antibodies (25). This may in part explain why variable results have been observed in clinical trials for convalescent plasma (25) compared to the relative efficacy of targeted antibody cocktails such as those marketed by Regeneron (26).

The probable involvement of M1 polarized macrophages in damaging surrounding tissues when non-neutralizing antibodies are present during infection with SARS-CoV-2 is problematic for many of the vaccines being developed for COVID-19, as most currently rely on production of antibodies using variations in delivery of the spike protein alongside an adjuvant to elicit production of an Th1 response. The preference for Th1-directed vaccines in COVID-19 originated due to historical observations of the association between Th2/Th17 responses in other virus models and subsequent immunopathology (27). Given what has been documented thus far for SARS-CoV-2 and the impossibility of preventing associations between macrophages and antibodies in the context of infection, this strategy may need to be reconsidered in favor of promoting a more balanced immune response. As yet, no antibody-dependent enhancement of disease has been observed in Phase I and II clinical studies (28) suggesting that despite production of non-neutralizing antibodies alongside those directed against the RBD there is little risk to vaccinated individuals as long as a sufficient titer of anti-RBD antibodies is generated and maintained.

However, once immunity starts to wane, a high ratio of non-neutralizing to neutralizing antibodies alongside M1 polarization, may be riskier than RBD-specific vaccination and may suggest that neutralizing antibody titers in vaccinated individuals should be monitored regularly to establish timelines for administration of booster doses. Lastly, the impact of M1 observed here and in the literature suggests that recalibration of COVID-19 vaccines to produce a more balanced Th1/Th2 response may alleviate side effects, leading to greater compliance from vaccine-hesitant populations, without sacrificing efficacy against disease.

## Materials and Methods

### Macrophage Infection Assays

Low passage Raw264.7 cells (ATCC TIB-71) were cultured in DMEM containing, 4 mM L-glutamine, sodium pyruvate (Hyclone SH30243), penicillin/streptomycin (Gibco 15140-122) and 10% FBS (Gibco 16000-044). The day prior to the assay, cells were seeded at a density of 1 x 10^4^ cells in 96 well tissue culture plates (Costar 3904) and incubated overnight at 37°C with 5% CO2. The following day cells were washed twice with 200ul of 1XPBS (Hyclone SH30256) and following the second wash, allowed to incubate in 100 ul of DMEM containing penicillin/streptomycin and 5% FBS (VIM).

Prior to the addition of virus, serial dilutions of antibodies against either the nucleocapsid (Genetex GTX632269) or spike proteins (polyclonal anti-spike, Abcam ab272504; anti-RBD, AcroBiosystems SAD-S35-100ug) of SARS-CoV-2 in a total volume of 60ul of VIM were made in a separate 96 well dilution plate to which 60 pfu/well of SARS-CoV-2 (USA-WA1/2020) was added for a final MOI of 0.05. The dilution plate was then incubated for 1 hour at 37°C, 5% CO2. Following incubation, media on the cells was replaced with 100 ul of the antibody and virus mixture and incubated for two days at 37°C, 5% CO2.

Supernatants were removed and stored at -80 °C for subsequent experiments with Vero E6 cells. Media was replaced with 100ul of VIM and viability of macrophages was then assessed by the addition of 100ul of Cell Titer Glo (Promega G7573) per well which was read for luminescence (Biotek) following incubation at room temperature for 5 minutes.

### Determination of Viral Load

Raw 264.7 cells were seeded at a density of 2×10^6^ cells/well in a 6-well plate (Geiner Bio-One 657165) and allowed to settle overnight at 37°C, 5% CO2. The following day, cells were infected with SARS-CoV2 with an MOI of 0.05 and incubated overnight at 37 °C with 5% CO2. Cells were lysed with 1 ml Trizol® (Ambion 15596018) and RNA extracted using a Direct-zol™-96 MagBead RNA miniprep kit (Zymo R2102). Quantitative PCR was performed using the VIRSeek SARS-CoV2 assay (Eurofins Genescan Technologies, Freiberg, Germany).

### Activation of MPs with LPS

*Escherichia coli* (Microbiologics 0617K) was grown overnight in BHI (Neogen NCM0016A) at 37°C. The culture was then diluted with 1X PBS to an OD of 1.0 in BAX System Lysis Buffer (Hygiena ASY2011) and heated for 10 minutes at 95 °C. Sterility was confirmed by inoculation of 20 ul of the resulting lysate into DMEM, which was then incubated at 37 °C for 24 hours prior to use in the macrophage assay at a volume of 10 ul per well for a 96 well plate or 30 ul per well for a 6 well plate. Negative controls received BAX System Lysis buffer without LPS at an equivalent volume.

### Evaluation of ACE2 Receptor Expression in Raw264.7 cells

Raw 264.7 cells were seeded onto sterile glass coverslips (VWR 16004-304) in a 6-well plate (Geiner Bio-One 657165) at a density of 2×10^6^ cells/well and allowed to settle overnight. Polyclonal anti-spike at a concentration of 6.25 ug/ml (described above) and/or LPS were added as appropriate in 1.5 ml of VIM which replaced the previous media after which the plate was returned to the incubator. The following morning, cells were infected with SARS-CoV2 with an MOI of 0.05 and incubated overnight at 37 °C with 5% CO2. Following incubation, all media was removed and the cells were fixed for 24h with 4% paraformaldehyde.

After fixation, coverslips were washed twice with 1X PBS for 5 minutes at room temperature prior to the addition of 2 mls of 1XPBS containing 0.5% Triton X-100 (Sigma X100-1L) and 5% w/v nonfat dry milk. Coverslips were incubated overnight at 4 °C in blocking buffer and probed the following morning with 1 :500 rabbit anti-ACE2 (Abcam ab15348) for 1h at room temperature on an orbital rocker. Following incubation, samples were washed three times with 1XPBS+0.2% Triton-X100 for 5 minutes, prior to incubation with 1:1,000 anti-rabbit IgG secondary antibody (Rockland 611-141-122) for 1 hour. Following incubation, samples were once again washed three times with 1XPBS+0.2% Triton-X100 for 5 minutes and mounted onto slides with ProLong Gold antifade reading with DAPI (Invitrogen P36935). Images were captured using a Zeiss LSM 710 confocal microscope and receptor expression quantitated using ImageJ.

### Impact of MP Supernatants on Vero E6 Susceptibility to SARS-CoV-2

Vero E6 cells (ATCC VERO C1008) were cultured in DMEM containing, 4 mM L-glutamine, sodium pyruvate (Hyclone SH30243), penicillin/streptomycin (Gibco 15140-122) and 10% FBS (Gibco 16000-044). The day prior to the assay, cells were seeded at a density of 1 x 10^4^ cells in 96 well tissue culture plates (Costar 3904) and allowed to incubate overnight at 37°C with 5% CO2. The following day cells were washed twice as described above with PBS and 100ul of VIM added to each well.

Prior to the addition of virus, 20 ul of the appropriate macrophage supernatant or DMEM was added to 40 ul of VIM for a total volume of 60ul in a separate 96 well dilution plate. SARS-CoV-2 (USA-WA1/2020) was added for a final MOI of 0.05 (60pfu/well in a volume of 60 ul) and 100 ul of the resulting supernatant+virus mixture was then added to the Vero E6 plate by replacing the media. The plate was then incubated for 3 days at 37°C with 5% CO2 and viability of the cells assessed as described for the macrophage assay described above.

### Quantitation of Inflammatory Mediators in Macrophage Supernatants

To evaluate the presence of cytokines and chemokines in the supernatants of SARS-CoV-2 infected macrophages, 50 ul of each supernatant was evaluated in triplicate using the Mouse Cytokine & Chemokine 36-Plex Procarta 1A Panel (ThermoFisher EPXR360-26092-901) as per manufacturer’s instructions. Data was collected using a Luminex FlexMap3D.

### Statistical Analysis

All graphical presentations of data and ANOVA analysis was conducted in GraphPad Prism 9.

## Presentation Statement

Preliminary data from this article was presented at IDWeek 2020 (*Open Forum Infectious Diseases*, Volume 7, Issue Supplement_1, October 2020, Page S319, https://doi.org/10.1093/ofid/ofaa439.701)

